# Impact of the aryl hydrocarbon receptor on Aurora A kinase and the G2/M phase pathway in hematopoietic stem and progenitor cells

**DOI:** 10.1101/2022.05.27.492132

**Authors:** Anthony M. Franchini, Keegan Vaughan, Soumyaroop Bhattacharya, Kameshwar Singh, Thomas A. Gasiewicz, B. Paige Lawrence

## Abstract

Recent evidence suggests that the environment-sensing transcription factor aryl hydrocarbon receptor (AHR) is an important regulator of hematopoiesis. Yet, the mechanisms and extent of AHR-mediated regulation within the most primitive hematopoietic cells, hematopoietic stem and progenitor cells (HSPCs), are poorly understood. Through a combination of transcriptomic and flow cytometric approaches, this study provides new insight into how the AHR influences HSPCs. Comparative analysis of intraphenotypic transcriptomes of hematopoietic stem cells (HSCs) and multipotent progenitor (MPP) cells from AHR knockout (AHR KO) and wild-type (WT) mice revealed significant differences in gene expression patterns. Notable among these were differences in expression of cell cycle regulators, specifically an enrichment of G2/M checkpoint genes when *Ahr* was absent. This included the regulator Aurora A kinase (*Aurka*, AurA). Interrogation of AurA protein levels in HSPC subsets using flow cytometry, in combination with inducible AHR KO or *in vivo* AHR antagonism showed that attenuation of AHR increased levels of AurA in HSCs and lineage-biased MPP cells. Overall, these data highlight a potential novel mechanism by which AHR controls HSC homeostasis and HSPC differentiation. These findings advance the understanding of how AHR influences and regulates primitive hematopoiesis.

**Highlights (max 85 characters):**

- AHR alters gene expression during HSC-MPP transition.
- Transcriptomic analysis shows AHR regulation of key G2/M phase regulators
- Inducible AHR KO mice show increased AurA levels in HSPC populations
- Acute antagonism of AHR increased AurA levels across multiple HSPC populations

## Introduction

Self-renewing hematopoietic stem cells (HSCs) give rise to all lineages of blood and immune cells [1]. HSCs remain predominantly quiescent during an organism’s lifespan, proliferating at key moments to replenish lineage committed progenitors, which give rise to mature cells [2]. In addition to HSCs, lineage-restricted multipotent progenitor cells (MPPs) contribute to primitive stages of hematopoiesis, and maintain downstream differentiated progenitors of all blood and immune cell populations [3, 4]. Collectively, HSCs and MPPs are referred to as hematopoietic stem and progenitor cells (HSPCs). HSPCs are dependent on external signals that maintain self-renewal, prompt expansion, and influence cell programming in order to restock progenitors and downstream cell lineages at steady state and in response to stressors, such as infection or malignancy [5]. Disrupted regulation of the balance between HSPC dormancy and proliferation can have serious long-term consequences [6, 7]. Also, maintaining stem and progenitor cell quiescence prevents premature stem cell exhaustion [8]. Yet, a critical function of HSPCs is rapid response to environmental cues, and part of this involves responding to external signals that modulate HSCs dormancy vs HSC proliferation and self-renewal, and the transition of HSCs into MPPs. Signals that regulate the balance between these cellular states are mediated by numerous cytokines, growth factors, and transcription factors [9].

One such regulator, the aryl hydrocarbon receptor (AHR), has been linked to alterations of immune cells and myeloproliferative disorders [10-14]. The AHR is a member of the Per-ARNT-Sim family, which encompasses a range of environment-sensing transcriptional regulatory factors [15]. Although experimental evidence supports a role of AHR in hematopoiesis [11, 16-18], large gaps in knowledge remain. For example, prior studies showed changes in functional status and frequency of HSCs and lineage committed hematopoietic cells in the absence of the AHR [13, 16, 18-20]. This includes the observation that the loss of AHR impacts lineage potential of HSPCs, including biasing towards myeloid-biased lineage precursors [19]. In other studies, antagonism of the AHR in human HSCs supports that it plays a role in regulating HSC proliferation and differentiation to progenitor cells [21]. However, the genes influenced by AHR in the context of regulating HSCs and MPP cells (i.e., HSPCs) have yet to be fully elucidated.

The study presented here was undertaken to better define the role of AHR in regulating HSPCs, and to identify genes and pathways influenced by AHR during the earliest stages of hematopoiesis. Using a combination of transcriptional and flow cytometric approaches, we present evidence that AHR shapes the HSPC transcriptome and differentiation program, influencing processes and pathways involved in critical checkpoints of cell cycle, including Aurora A kinase. These observations expand the role of AHR as a central regulator of primitive stages of hematopoiesis.

## Materials and Methods

### Mice and *in vivo* treatment

C57BL/6J wild-type (WT) mice were purchased from the Jackson Laboratory (Bar Harbor, ME). Initial breeding stocks of B6.129-Ahr^tm1Bra^/J (AHR KO) and Ahr^tm3.1Bra^/J (AHR^Fx/Fx^) mice were provided by Christopher Bradfield (University of Wisconsin, Madison, WI) [22, 23]. B6.129-Gt(ROSA)26Sortm1(Cre/ERT2)Tyj/J (CRE^ERT2^) mice were purchased from the Jackson Laboratory [23, 24] and crossed with AHR^Fx/Fx^ mice to generate AHR^Fx/Fx^CRE^ERT2^ mice [19]. All data presented are from female mice that were 6–10 weeks of age at the time of experiments. Excision of *Ahr* from tamoxifen-treated AHR^Fx/Fx^ CRE^ERT2^ (AHR iKO) mice was confirmed by PCR [19, 22]. All primers used for genotyping PCR can be found in supplemental table S1.

To assess *in vivo* proliferation, mice were administered 120 mg bromodeoxyuridine (BrdU, Sigma-Aldrich, St. Louis, MO) per kg body weight by intraperitoneal (i.p.) injection 2h prior to the termination of the experiment. AHR^Fx/Fx^ and AHR^Fx/Fx^CRE^ERT2^ mice were administered 25 mg tamoxifen/kg body weight by i.p. injection on 3 consecutive days [19, 25]. After the third dose of tamoxifen, mice were allowed to rest for 14 days prior to assessment. Tamoxifen was purchased from Sigma-Aldrich (St. Louis, MO). In some studies, mice were given a single i.p. injection of 150 µg 5-fluorouracil (5-FU, Sigma-Aldrich) per kg of body weight [26, 27]. For AHR antagonism, CH-223191 (Sigma-Aldrich) was dissolved in corn oil at a concentration of 0.5 mg/ml, and 100 μg per mouse was delivered by i.p. injection [13, 28, 29]. All animal treatments had prior approval of the Institutional Animal Care and Use Committee of the University of Rochester. The University is accredited by the Association for Assessment and Accreditation of Laboratory Animal Care (AAALAC). Animals were treated humanely and with due consideration to alleviation of distress and discomfort, following U.S. Public Health Service Policy on Human Care and Use of Laboratory Animals guidelines for the handling of vertebrate animals.

### Collection of bone marrow cells

Both femurs and tibias were excised from each mouse, cleared of adherent tissue, and crushed in mortar and pestle to release the bone marrow cells [11, 16, 18, 30]. Bone marrow cells were resuspended in Iscove’s Modified Dulbecco’s Medium (IMDM, Gibco, Gaithersburg, MD) supplemented with 2.5% fetal bovine serum (HyClone, Logan, UT). Erythrocytes were lysed using an ammonium chloride solution (0.15M NH_4_Cl, 10mM NaHCl_2_, 1 mM Na_2_EDTA) for five minutes at room temperature, and bone marrow cells were passed through 40 µm filters twice to remove debris. Cells were immediately used for analysis by flow cytometry, or subjected to further purification and extraction of cellular materials.

### HSPC isolation and flow cytometry

For cell sorting prior to RNA isolation, lineage positive (CD5, CD45R, CD11b, Gr-1, 7-4, Ter-119) bone marrow cells were removed by positive selection using immunomagnetic beads (Miltenyi, Waltham, MA) prior to labeling remaining cells for fluorescence activated cell sorting (FACS). Lineage negative cells were incubated with anti-CD16/32 to block non-specific binding, and labeled with antibodies against Sca1, cKit, CD135, CD48, CD150, as well as a cocktail of lineage-specific antibodies to identify MPPs (Lin^-^CD135^-^ckit^+^Sca1^+^CD48^+^CD150^+/-^) and HSCs (Lin^-^CD135^-^ckit^+^Sca1^+^CD48^-^CD150^+^). Sorting of HSC and MPP populations was performed using a BD FACS Aria II flow cytometer (BD Biosciences, San Jose, CA) in the University of Rochester Flow Cytometry Core. Details regarding the antibodies used are provided in supplemental table S2. The gating strategy for sorting HSCs and MPPs is presented in supplemental figure 1.

To identify distinct HSPC populations using analytical flow cytometry, bone marrow cells were incubated with fluorescently tagged antibodies that recognize the following cell surface markers: CD117, CD48, CD34, CD135, CD48, CD150 and lineage markers (CD3, CD45R, CD11b, Ly6G, and TER119) [31]. Non-specific binding was blocked by pre-incubation of cells with anti-CD16/32. After cell surface labeling, cells were fixed using 2% paraformaldehyde and analyzed directly by flow cytometry, or permeabilized using 1% saponin (Sigma-Aldrich) to detect Aurora A kinase. Intracellular Aurora A kinase (AurA) was assessed using a polyclonal antibody (ST46-07, Novus Biologicals, Littleton, CO), in combination with fluorescently-labeled donkey anti-rabbit IgG antibody (Biolegend). For measuring BrdU incorporation, cells were permeabilized with Triton X-100 (0.5%, Sigma-Aldrich) and a directly conjugated anti-BrdU monoclonal antibody was used. A list of all antibodies, including fluorochrome, vendor, and amount used is in supplemental table S2. Fluorescence minus one (FMO) controls were used to determine non-specific fluorescence and define all gating parameters [32]. Two to three million bone marrow cells from individual mice were stained, and 1 million events were collected using a LSRII flow cytometer (BD Biosciences, San Jose, CA). Data were analyzed using the FlowJo software program (TreeStar, Ashland, OR). The specific combinations of molecular markers used to identify different HSPCs are denoted in the figure legends. The gating strategy used to identify HSPC subsets (i.e., HSCs, MPP1, MPP2, MPP3 and MPP4 cells) is presented in supplemental figure 2.

### RNA Sequencing library construction and transcriptomic analysis

RNA was isolated from sorted HSCs and MPPs using RNeasy Mini kits (Qiagen, Valencia, CA). The concentration of RNA was determined using an Agilent 2100 bioanalyzer (Agilent Technologies). RNA (1 ng) was amplified using SMARTer library Ultra Low cDNA v4 kit (Takara Bio, Mountainview, CA). Sequencing cDNA libraries were preparing using a NexteraXT DNA library prep kit (Illumina, San Diego, CA). The cDNA library (150 pg/sample) was sequenced on an Illumina HiSeq2500 system to generate ∼20 million 100-bp single end reads per sample. Sequences were aligned against the mouse genome mm10 using the Splice Transcript Alignment to a Reference (STAR) algorithm [33], counted with HTSeq [34], and normalized for total counts (counts per million, CPM) A non-specific filtering strategy was used to remove genes with low expression values. Samples were excluded based on poor read count/mapped read numbers, or if shown to be extreme outliers by hierarchical clustering and principal component analysis (PCA). Differential gene expression was assessed by paired DESeq2 [35] to identify genes with significant differences in mean expression (false discovery rate, FDR, < 0.05). Differentially expressed genes (DEGs) identified by direct comparison of datasets from HSCs (across genotype) and MPPs can be found in supplemental tables S3 and S4, respectively. DEGs identified by intraphenotypic analysis (i.e., HSC vs. MPP within genotype) are shown in supplemental table S5 and S6, respectively.

### Statistical analyses

With the exception of sequencing data, statistical analyses were performed using JMP Pro 14.0.0 (SAS Institute, Cary, NC). Differences between means of multiple independent variables were compared using one-way ANOVA followed by post hoc tests (Tukey honestly significant difference or Dunnett’s test). Differences between two groups at a single point in time were analyzed using Student’s t test. The slope of the line was calculated using goodness of fit modeling and derivation from non-zero slope determination was performed. Differences in mean values were considered statistically significant when p-values were <0.05. Error bars on all graphs represent the SEM. Linear regression was performed using Prism (Version 8, GraphPad). R^2^ and p-value of the correlation were calculated using a goodness of fit model and compared to a non-zero slope are shown for each regression analysis.

### Data Availability

RNA-sequencing data have been deposited to the NIH Gene Expression Omnibus (GEO) under accession number (GSE163284).

## Results

### Absence of AHR alters HSPC transcriptome

Accumulating evidence suggests that AHR is necessary for the development and function of the mammalian immune system [36, 37]. Consistent with previous studies [16, 19, 20], isolated bone marrow cells from AHR KO mice have a greater frequency of HSCs compared to bone marrow cells from WT mice (Figure 1A). We show here that there was a 28% increase in BrdU+ HSCs in AHR KO mice compared to HSCs from WT mice (Figure 1B). The proliferative capacity of HSPCs is integrally linked with their transcriptional programming and differentiation [9, 38, 39]. However, the full extent of the gene network regulated by AHR in HSCs and MPP cells is incomplete as prior studies relied upon microarray analysis or lacked markers to refine differences between HSPC subsets [18, 20]. To directly address this, we evaluated the transcriptome of HSCs and MPPs in the absence and presence of AHR. Specifically, HSCs and MPPs from WT and AHR KO mice were isolated using FACS followed by high throughput RNA sequencing (RNA-Seq; Figure 1C). Unsupervised hierarchical clustering confirmed delineation of HSCs and MPPs into separate clusters (Figure 1D). Furthermore, within HSPCs, HSCs and MPPs clustered within the same genotype (Figure 1D). Comparison of gene expression profiles in HSCs from WT and AHR KO mice revealed 103 differentially expressed genes (DEGs; Figure 1E, and supplemental table S3). In contrast, comparison of expressed genes in MPPs from WT versus KO mice showed only six DEGs. Among these, only *Garnl3* was shared with DEGs from HSCs, and the other 5 were distinct from the DEGs in HSCs (Figure 1F, and supplemental table S3). Among these six DEGs in MPPs, two encode receptors (*Ffar2* and *Slamf6*), two encode proteins involved cell metabolism (*Akr1c13, Tmem18*), two are for factors involved in cell signaling (*Garnl3, Mrvi1*, supplemental table S4).

**Figure 1.**
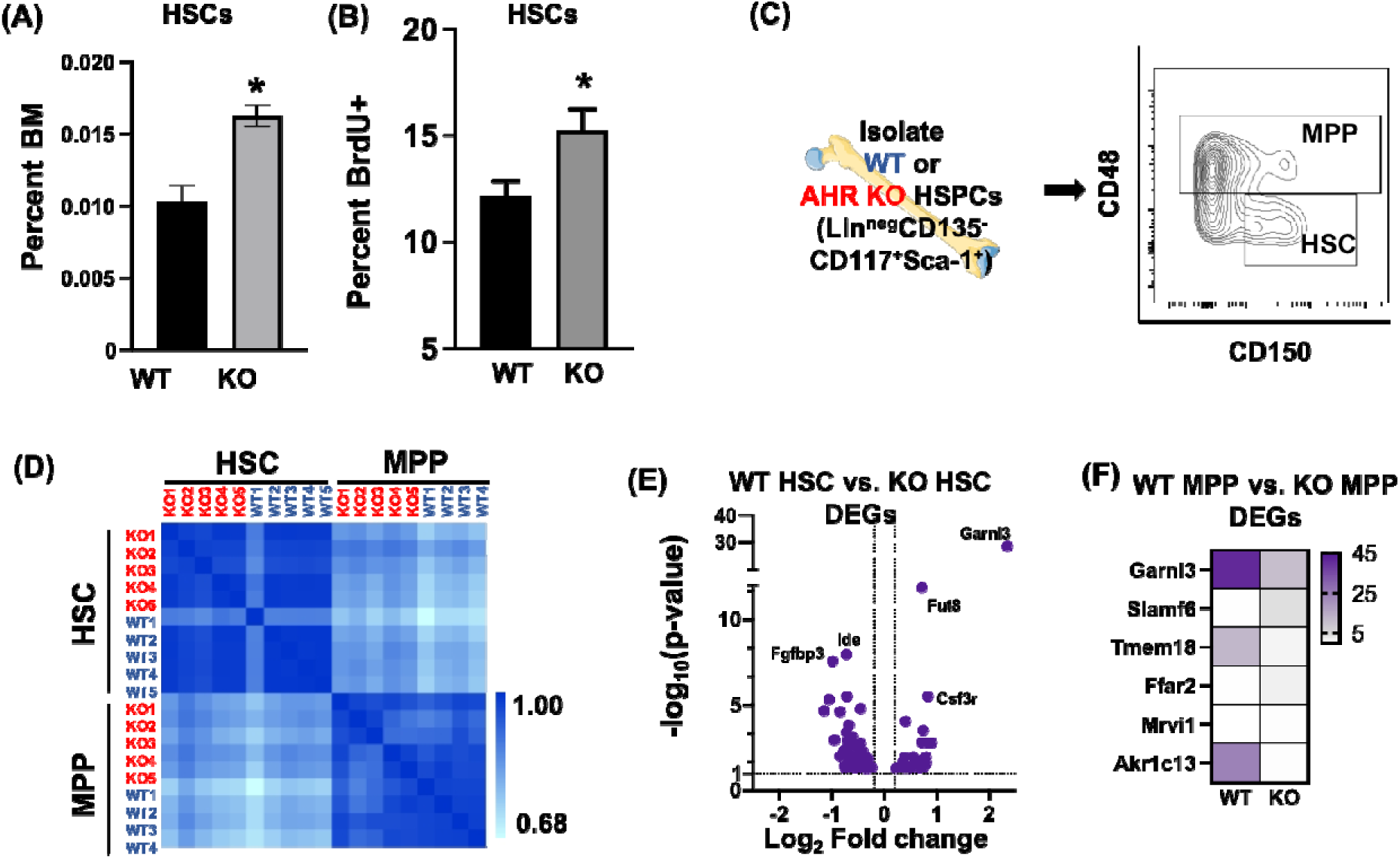
Gene expression patterns in HSC and MPP from wild-type and AHR KO mice. The percentage of (A) HSCs and (B) BrdU+ HSCs, measured by intracellular flow cytometry staining (n=5) from WT and AHR KO mice was determined by flow cytometry (n=5 mice per genotype). HSC were defined as Lin^-^CD135^-^ CD117^+^Sca1^+^CD48^-^CD150^+^ cells. BrdU was delivered i.p. 2 h prior to bone marrow cell isolation. Asterisks denote p<0.05 between genotype, Student’s t-test. (C) HSCs and MPPs from individual C57BL/6 (WT) mice and AHR KO mice were isolated by cell sorting (FACS) followed by RNA-Seq. HSCs were defined as Lin^-^ CD135^-^CD117^+^Sca1^+^CD48^-^CD150^+^ cells, and MPPs were defined as Lin^-^CD135^-^CD117^+^Sca1^+^CD48^+^CD150^+^ cells. There were five independent biological replicates of HSCs and MPPs per genotype. (D) Unsupervised hierarchical clustering of RNA-Seq datasets was performed. (E) Volcano plot depicts the 103 DEGs identified between the WT and AHR KO HSCs datasets. See supplemental table S3 for the complete list of these DEGs, with fold change and p-values. (F) Heatmap depicts the six genes that were differentially expressed in MPPs from WT and AHR KO mice. See supplemental table S4 for fold change and p-values for each DEG.

At first glance, this transcriptomic assessment did not offer particularly compelling insight into the regulatory role of AHR in HSPCs. To further understand how AHR influences gene expression in HSPCs, we compared transcriptomic profiles in HSCs vs MPPs within and across genotypes. In total, 1586 DEGs were identified in the HSC to MPP intraphenotypic dataset from WT mice (Figure 2A, supplemental table S5). In the intraphenotypic dataset from AHR KO mice, there were 2474 DEGs (Figure 2A, supplemental table S6). Comparative analysis of these intraphenotypic datasets (i.e., comparing WT and AHR KO HSC-MPP DEG datasets) showed that 1124 DEGs were common to both genotypes. In contrast, 1315 DEGs were unique to the AHR KO HSC to MPP intraphenotypic dataset, while 460 genes were unique to the WT HSC to MPP dataset (Figure 2A). This two-way comparative approach indicates that absence of AHR has a more pronounced impact on HSPCs across the differentiation process (i.e., HSCs vs MPPs), rather than on a singular cellular state.

**Figure 2.**
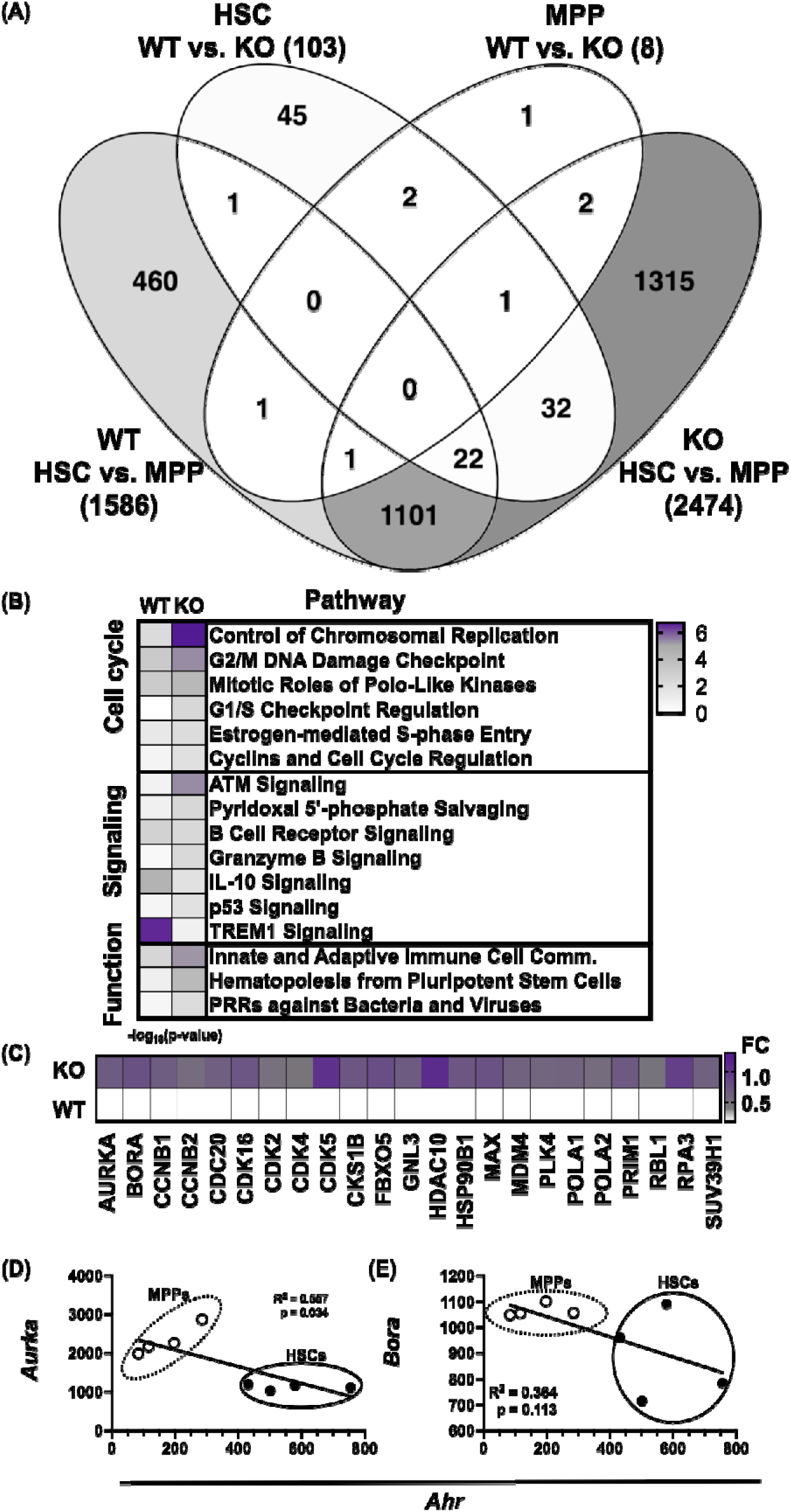
Absence of AHR alters cellular programming of HSPCs. (A) Four-way Venn diagram depicts the number of differentially expressed genes (DEGs) in the following comparisons: WT HSC vs WT MPP, KO HSC vs KO MPP, WT HSC vs KO HSC, and WT MPP vs KO MPP. The number in parentheses indicates the total number of DEGs each of these comparisons. The complete lists of DEGs from these comparisons are in supplemental tables S5 and S6. (B) Pathways identified as significantly different in the WT and KO intraphenotypic HSPC datasets (i.e., comparing DEGs in HSCs and MPPs across genotype). The heatmap denotes -log_10_(p value) in WT and KO intraphenotypic HSPC datasets (numerical data are in supplemental table S7 and S8). (C) Heatmap depicts the log_2_ fold change of cell cycle DEGs identified in the six cell cycle pathways shown in panel B. The fold change and p-value for each DEG can be found in supplemental table S4. (D-E) Linear regression analysis was performed on the normalized reads of *Ahr, Aurka, and Bora* using RNA-Seq data set described in Figure 1. R^2^ and p-values are shown on each graph. Solid (HSC) and open (MPP) circles denote data from individual samples. The numerical information represented in each plot is supplemental tables S3 and S4.

Pathways analysis was utilized to compare and relate changes to the transcriptional landscape of HSPCs in order to unbiasedly identify functional clusters or cellular pathways affected by the absence of AHR. Comparison of the DEGs in the WT and KO intraphenotypic datasets identified 37 unique pathways attributed to the lack of AHR. The most affected pathways, ranked by p-value, were related to cell cycle, signaling, and HSC function (Figure 2B, supplemental table S7 and S8). Within the six cell cycle pathways that were significantly different, 23 DEGs were unique to the AHR KO intraphenotypic dataset (Figure 2C). That is, these genes were not differentially expressed in the HSC-MPP comparison from WT mice, but were differentially expressed between HSCs and MPPs from AHR KO mice. Notable within the DEGs was the polo-like kinase and G2/M phase regulator Aurora A kinase (*Aurka*, AurA), which exhibited a 1.6-fold increase in gene expression (0.67 log_2_ fold change) in the KO dataset (Figure 2C). In addition to increased *Aurka*, absence of AHR correlated with greater expression of the gene for its dimerization partner, *Bora*, which was 1.7-fold-higher in the AHR KO data set, compared to WT dataset (Figure 2C, supplemental tables S3 and S4). Furthermore, within the WT HSC-WT MPP datasets, there was a statistically significant inverse correlation between *Aurka* and *Ahr* levels (Figure 2D). Specifically, MPPs expressed higher levels of *Aurka* and lower *Ahr* levels, while HSCs possess lower *Aurka* and *Ahr* levels, respectively. *Bora* expression levels showed a similar pattern, but *Bora* expression in HSCs was more variable than *Aurka*, therefore there was not a significant correlation with Ahr expression levels (Figure 2E). These data are consistent with the idea that not all cell cycle pathways are regulated by the same mechanism [11, 40-43]. Furthermore, these observations suggest that AHR exerts some level of regulatory control over HSPC function via its control of the G2/M phase checkpoint.

### Acute loss or antagonism of AHR results in increased AurA in HSPCs

To further assess the connection between AHR and Aurora A kinase in HSPCs, relative levels of AurA protein were examined using flow cytometry. This approach has a distinct advantage over immunoblotting in that AurA protein in HSPCs, as well as within phenotypically distinct HSPC subpopulations, can be examined without pooling cells from large numbers of mice. Consistent with its ubiquitous expression in other cell types [44], AurA was detected in over 99% of all HSPCs (Figure 3A). HSPCs include HSCs and MPPs, and MPPs include four sub-populations (MPP1 through MPP4 cells), which can be distinguished using well defined cell surface markers [31] (supplemental figure 2). Essentially all of these HSPC subpopulations expressed AurA, with no discernable difference in the percentage of AurA+ cells among HSCs or MPPs (Figure 3B). Furthermore, the mean fluorescence intensity (MFI) of AurA was not significantly different among all of the HSPC subsets, (Figure 3C) indicating similar levels of expression amongst these different sub-populations.

**Figure 3.**
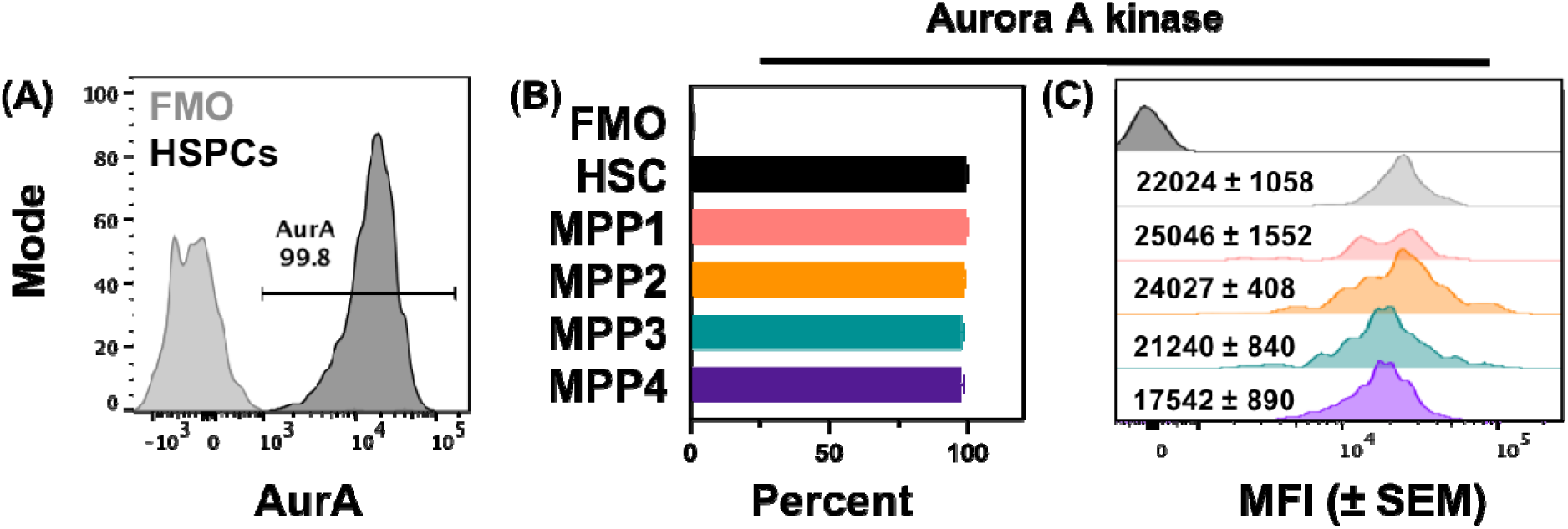
Aurora A kinase protein levels are similar in HSPC subsets. Bone marrow cells from C57Bl/6 mice (n=5) were isolated and stained for flow cytometry. (A). The dark grey histogram depicts AurA expression in HSPCs. The fluorescence minus one (FMO) control is shown in light grey. (B) Graph depicts the percentage of different HSPC subsets that were AurA+. There were five mice used, and bone marrow cells from individual mice were not pooled. Error bars denote SEM. Subsets were defined as follows: HSC (Lin^-^ Sca1^high^ cKit^+^ CD34^-^ CD135^-^ CD48^-^ CD150^+^), MPP1 (Lin^-^ Sca1^high^ cKit^+^ CD34^+^ CD135^-^ CD48^-^ CD150^+^), MPP2 (Lin^-^ Sca1^high^ cKit^+^ CD34^+^ CD135^-^ CD48^+^ CD150^+^), MPP3 (Lin^-^ Sca1^high^ cKit^+^ CD34^+^ CD135^-^ CD48^+^ CD150^-^), and MPP4 cells (Lin^-^ Sca1^high^ cKit^+^ CD34^+^ CD135^+^ CD48^-^ CD150^-^). (C) Histograms represent the mean fluorescent intensity (MFI) of AurA in each HSPC subset. The number indicates the MFI (±SEM). A detailed gating strategy is provided in supplemental figure 2.

To determine if the absence of AHR alters AurA levels in HSPCs, AHR^Fx/Fx^ mice were crossed with Cre^ERT2^ mice, to generate stable lineages of AHR^Fx/Fx^Cre^ERT2^ mice, which upon treatment with tamoxifen, are inducible AHR knockout (AHR iKO) mice [19]. Two weeks after tamoxifen treatment, *Ahr* was fully excised in bone marrow cells from AHR iKO mice, whereas *Ahr* was not excised in tamoxifen-treated AHR^Fx/Fx^ mice (Figure 4A). The level of AurA was 20% higher in HSPCs from AHR iKO mice compared to HSPCs from AHR^Fx/Fx^ controls (Figure 4B-C). Within HSPC sub-populations, the greatest differences in AurA levels were in MPP1 and MPP3 cells (Figure 4D-H). Higher levels of AurA were also observed in HSCs, MPP2, and MPP4 subsets from iAHR KO mice, but mean values were not statistically different from AHR^Fx/Fx^ controls. Nonetheless, these findings indicate that acute AHR deletion was sufficient to alter the level of AurA within HSPCs.

**Figure 4.**
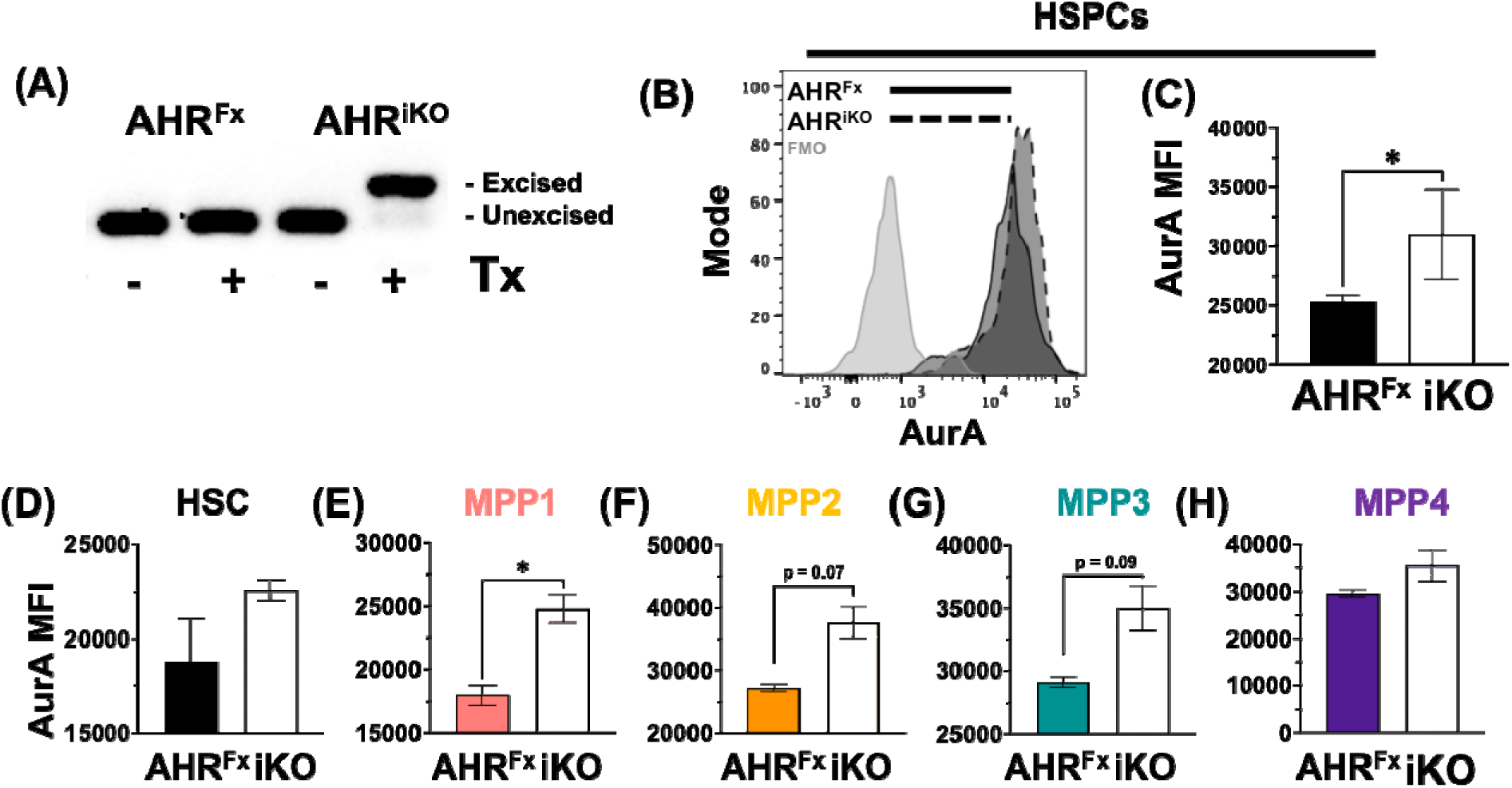
Inducible deletion of AHR increases AurA expression in HSPCs. AHR^Fx/Fx^ and AHR^CreERT2^ mice (5 mice per genotype) were administered tamoxifen i.p., 25 mg/Kg body weight (BW) daily for 3 consecutive days. Two weeks after final tamoxifen (Tx) treatment, bone marrow cells from AHR^Fx/Fx^ and inducible AHR knockout (AHR iKO; iKO) mice were isolated and *Ahr* gene excision assessed by PCR. (A) The gel image shows PCR results obtained from one AHR^Fx/Fx^ with and without Tx treatment, and one AHR^Fx/Fx^Cre^ERT2^ with and without Tx treatment (labeled iAHR KO). (B) Representative histogram depicts AurA staining in HSPCs (Lin-CD117+Sca-1+ cells) from AHR^Fx/Fx^ (solid line) and AHR iKO (dotted line) mice (all mice received Tx). The fluorescence minus one (FMO) control is shown in light grey. (C) The graph shows the AurA MFI in HSPCs from AHR^Fx/Fx^ (AHR^Fx^) and AHR iKO mice (iKO). (D-H) Graphs depict the mean AurA MFI in (D) HSCs, (E) MPP1, (F) MPP2, (G) MPP3 and (H) MPP4 cells. Cell subsets were defined as follows (gating strategy in supplemental figure 2): HSC (Lin^-^ Sca1^high^ cKit^+^ CD34^-^ CD135^-^ CD48^-^ CD150^+^), MPP1 (Lin^-^ Sca1^high^ cKit^+^ CD34^+^ CD135^-^ CD48^-^ CD150^+^), MPP2 (Lin^-^ Sca1^high^ cKit^+^ CD34^+^ CD135^-^ CD48^+^ CD150^+^), MPP3 (Lin^-^ Sca1^high^ cKit^+^ CD34^+^ CD135^-^ CD48^+^ CD150^-^) and MPP4 cells (Lin^-^ Sca1^high^ cKit^+^ CD34^+^ CD135^+^ CD48^-^ CD150^-^). The error bars indicate the SEM. Asterisks denote p<0.05 by Student’s t-test.

The AHR antagonist CH-223191 was used to further examine the relationship between attenuation of AHR and AurA levels. Wild-type mice were injected with CH-223191 or with the vehicle, corn oil (Figure 5). Administration of CH-223191 did not alter the frequency of HSPCs in the bone marrow (Figure 5A), nor did it affect the percentage of HSPCs that were AurA+ (Figure 5B). However, HSPCs from mice treated with CH-223191 had approximately 30% higher levels of AurA compared to HSPCs from mice given the vehicle control (Figure 5C). Similarly, AurA levels were 10-20% higher in HSCs, and MPP subsets following CH-223191 treatment compared to controls (Figure 5D-H). These data indicate that antagonism of AHR increased AurA levels within HSPC subpopulations, and that it is possible to modulate AurA levels by directly targeting AHR.

**Figure 5.**
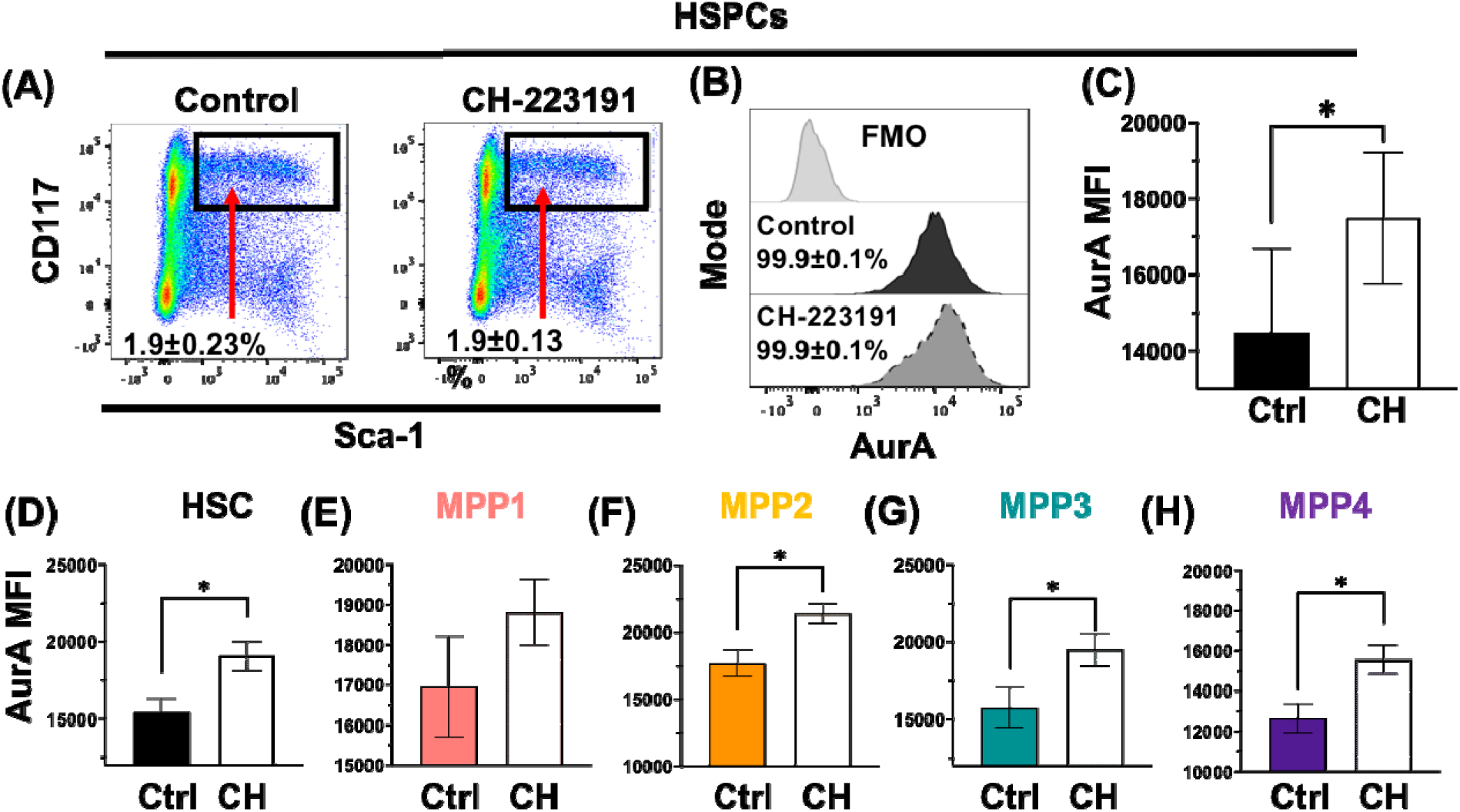
*In vivo* AHR antagonism enhances AurA expression in HSPCs. C57Bl/6 mice were administered a single dose of CH-223191 (100 µg; CH) or corn oil vehicle control (Ctrl) by i.p. injection (5 mice per group). Bone marrow cells were harvested 2 days later for analysis using flow cytometry. (A) FACS plots depict HSPCs (Lin^-^ CD117^+^Sca^-^1^+^) from control or CH treated mice. The number on each plot denotes the mean percentage of HSPCs (±SEM) in bone marrow. (B) Representative histograms depict AurA staining of HSPCs from Ctrl (black fill) and CH-treated mice (dark grey fill). The fluorescence minus one (FMO) control is shown in light grey. The numbers indicate the mean percentage (±SEM) of AurA-positive cells in each group. (C) Graph shows the MFI of AurA in HSPCs from Ctrl and CH treated mice. (D-H) Graphs depict the mean AurA MFI in (D) HSC, (E) MPP1, (F) MPP2, (G) MPP3 and (H) MPP4 cells. Cell subsets were defined as follows: HSC (Lin^-^ Sca1^high^ cKit^+^ CD34^-^ CD135^-^ CD48^-^ CD150^+^), MPP1 (Lin^-^ Sca1^high^ cKit^+^ CD34^+^ CD135^-^ CD48^-^ CD150^+^), MPP2 (Lin^-^ Sca1^high^ cKit^+^ CD34^+^ CD135^-^ CD48^+^ CD150^+^), MPP3 (Lin^-^ Sca1^high^ cKit^+^ CD34^+^ CD135^-^ CD48^+^ CD150^-^) and MPP4 cells (Lin^-^Sca1^high^ cKit^+^ CD34^+^ CD135^+^ CD48^-^ CD150^-^). The error bars indicate the SEM. Asterisks denote p<0.05 by Student’s t-test.

## Discussion

HSPCs are at the apex of hematopoiesis, and are responsible for providing all the cells of the blood and immune system across the entire lifetime. A key aspect of regulating HSPC function is balancing stem cell dormancy with the need to proliferate and differentiate into progenitor cells in response to external signals. This process is tightly controlled, and AHR has been put forth as an important regulator of hematopoiesis [14, 17-19]. The data presented in this study further this understanding, and expand the role of AHR as a regulator of cycle progression by providing evidence that the loss of AHR modulates expression levels of cyclin dependent kinases, polo-like kinases, and Aurora A kinase in HSPCs. Polo-like kinases, and specifically AurA, play critical roles in mitosis [45-47]. Cumulatively, the data presented in the current study support two interrelated ideas: that no singular AHR-gene interaction is solely responsible for differences in HSC quiescence-cell cycle entry, and that AHR is embedded in an integrated network of regulatory factors. The observation that loss or antagonism of AHR increased intracellular levels of AurA is novel and consequential. Increases in cell cycling reduce the future proliferative capacity of HSPCs, accelerate cellular senescence, and contribute to development of hematological diseases, such as myelodysplastic syndromes and leukemia [48-51]. Not only does absence of AHR correlate with a higher percentage of proliferating HSCs in mice (Fig 1), but inhibition of AHR directly promotes HSC proliferation *in vitro* [21]. Moreover, chronic myeloid leukemia cells express significantly lower amounts of *Ahr* transcript compared to healthy controls [52]. Therefore, loss or attenuation of AHR may remove or lessen the need for other signals to pass through this cell cycle checkpoint, resulting in aberrant HSPC proliferation.

Another key finding in this study was that the bulk of transcriptomic differences related to the absence or presence of AHR were uncovered when comparing gene expression differences between two different populations of cells, not by comparisons within the same cell type. These differences in gene expression were particularly notable in cell cycle pathways including the G1/S and G2/M phase checkpoint pathways, which further reinforces the idea that AHR is a regulator of cell cycle gene expression [39, 53-58]. While there were many DEGs in HSC versus MPPs, the limited number of DEGs detected when comparing HSCs in WT and AHR KO mice may reflect the fact that a large portion of the HSC pool is transcriptionally dormant [59]. The very limited number of DEGs in MPPs in WT versus AHR KO mice possibly reflect that heterogenous population that are MPPs [30]. Moreover, when considered in the context of the rich set of DEGs in the intraphenotypic datasets (ie, comparing DEGs in HSCs vs MPPs within and across genotype), suggests that AHR plays important role in regulation of the transition of HSCs into MPPs. HSCs and MPPs represent populations with distinct transcriptional profiles, and in the absence of AHR there is differential priming or responses of HSPCs to external stimuli. This differential response may explain the association of attenuation of AHR and myeloid lineage biasing of hematopoietic cells that has been reported [19, 20]. Yet, the current study suggests that AHR regulation of HSPCs and hematopoiesis is very complex, context specific, and that AHR may influence proliferaiton and differentiation via distinct mechanisms. By interacting with multiple cell cycle pathways, including AurA, it is also possible that AHR affects the balance between HSPC subsets, which is disrupted and leans towards the myeloid lineage when AHR is absent or its function is attenuated.

The ability to modulate HSPCs via AHR has the potential to alter the course of diseases. Hematopoiesis is a complex process wherein small alterations have significant downstream consequences on host defenses against infection, repair following injury, hematologic diseases, as well as tissue oxygenation and vascular integrity [5, 60, 61]. That the presence or absence of AHR affects gene expression of HSCs and MPPs indicates that it plays a role in the production of hematopoietic cells, and in part this may be via regulation of AurA. AurA is already a promising cancer therapy target (reviewed in [62]). Therefore, targeting the AHR-AurA and, more broadly, the AHR-polo-like kinase axis, provides an exciting new avenue for treating hematologic diseases. Given that AHR also affects proliferation of other cell types, it is possible that AHR-AurA connections contribute to AHR’s role regulating proliferation of non-hematopoietic cells in peripheral tissues, including the mammary gland, skin, and nervous systems [63-65]. Moreover, these new findings point to a broader role of AHR in other types of progenitor cells, such as those in the skin, liver and GI tract, which have high levels of AHR, are sensitive to modulation by AHR ligands, and in which AHR signaling has been associated with altered proliferation [59, 66-68]. Also, depending on context, exogenously provided AHR ligands can exacerbate or attenuate disease by modulating cell proliferation and differentiation [66, 69, 70]. That AHR can promote or dampen cellular processes may seem paradoxical, yet when properly contextualized with the transcriptomic findings presented herein, it suggests AHR helps cells sense and respond to external cues and integrate their responses within the complex regulatory networks that control cellular function. From this perspective, this current work furthers our appreciation of the complexity of the AHR as a conduit for environmental cues that modulate HSPCs, and provides insight into how AHR may affect other cells throughout the body.

## Acknowledgements

The authors thank Dr. John Ashton and the staff at the University of Rochester (UR) Genomics Research Center, and Dr. Timothy Bushnell, Matthew Cochran, and the staff of the UR Flow Cytometry Core for assistance in with cell sorting and RNA sequencing. This work was supported by grants from the National Institutes of Health (R01ES023260, R01ES004862, P30ES01247, and T32ES007026).

## Supplemental Figures and data

**Supplemental Figure 1.**
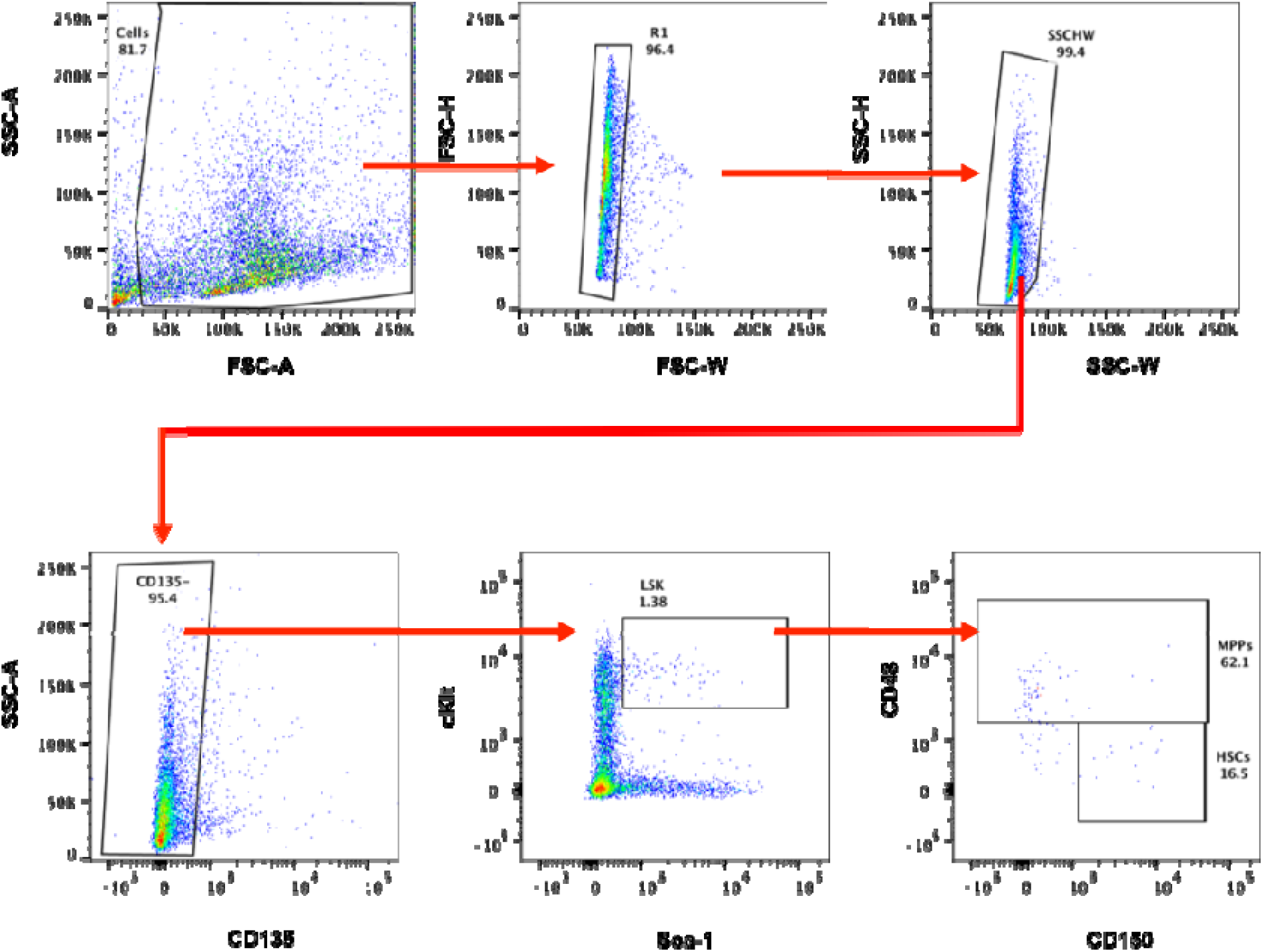
Flow cytometry gating used in isolation of HSCs and MPPs for RNA-Sequencing. Lineage depleted bone marrow cells were further isolated and purified by fluorescence-activated single cell sorting. Flow cytometry gating strategy used in the sorting of HSC and MPP subsets for RNA sequencing. HSCs were defined as Lin^-^CD135^-^CD117^+^Sca1^+^CD48^-^CD150^+^ cells, and MPP were defined as Lin^-^CD135^-^ CD117^+^Sca1^+^CD48^+^CD150^+^ cells.

**Supplemental Figure 2.**
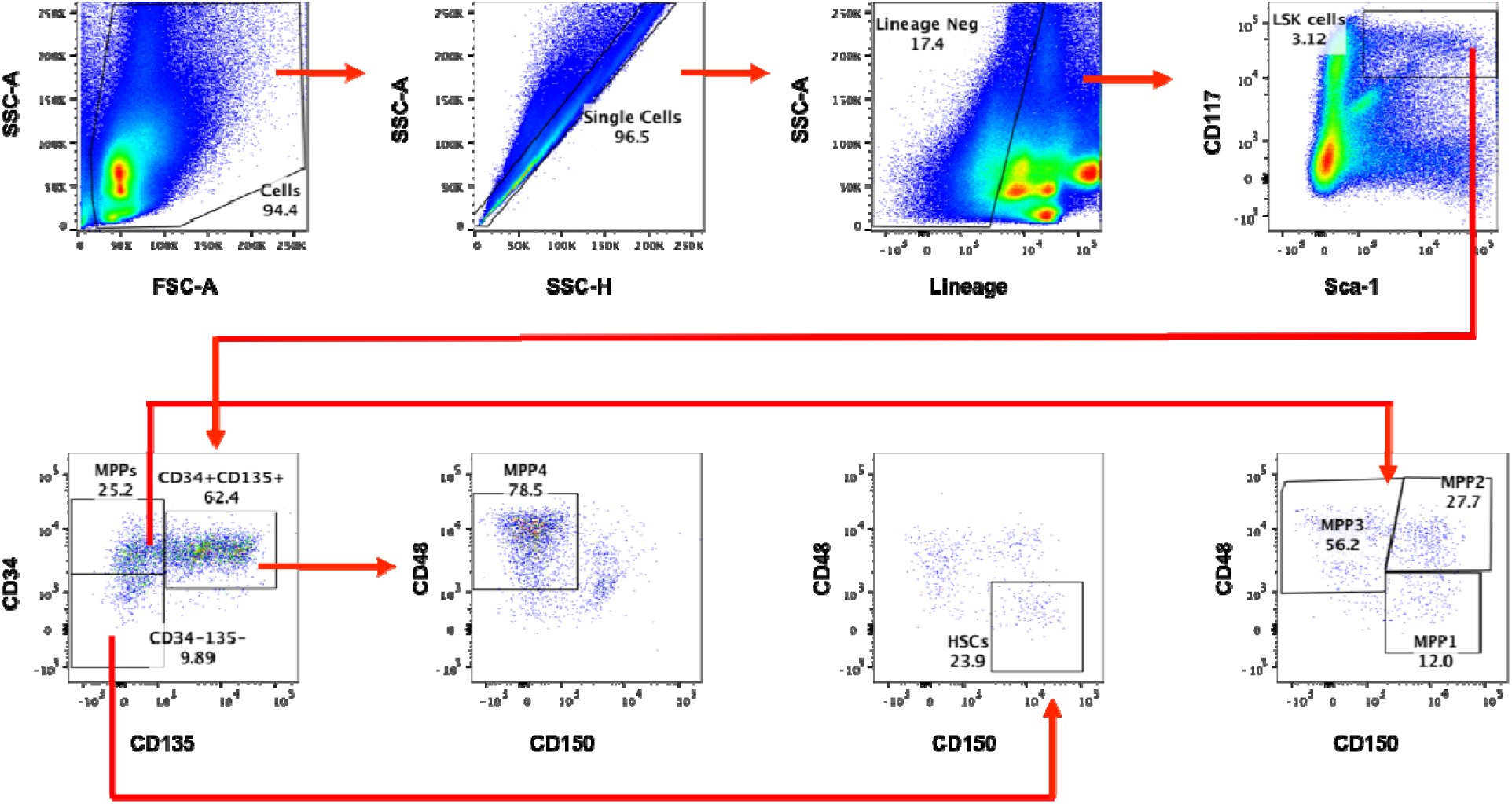
Flow cytometry gating strategy to measure AurA levels in HSPCs. Gating strategy use to identify HSPC subsets using flow cytometry. Cell subsets were defined as follows: HSCs (Lin^-^ Sca1^high^ cKit^+^ CD34^-^ CD135^-^ CD48^-^ CD150^+^), MPP1 (Lin^-^ Sca1^high^ cKit^+^ CD34^+^ CD135^-^ CD48^-^ CD150^+^), MPP2 (Lin^-^ Sca1^high^ cKit^+^ CD34^+^ CD135^-^ CD48^+^ CD150^+^), MPP3 (Lin^-^ Sca1^high^ cKit^+^ CD34^+^ CD135^-^ CD48^+^ CD150^-^) and MPP4 cells (Lin^-^ Sca1^high^ cKit^+^ CD34^+^ CD135^+^ CD48^-^ CD150^-^). Lin-denotes cells that do not express the following cell surface markers: CD3, CD45R, CD11b, Ly6G, and TER119.

**Table S1**. PCR primers for genotyping.

**Table S2**. Hematopoietic stem and progenitor cell staining panel used in flow cytometry.

**Table S3**. Differentially expressed genes identified between WT HSC and AHR KO HSCs.

**Table S4**. Differentially expressed genes identified between WT MPP and AHR KO MPP cells.

**Table S5**. Differentially expressed genes identified by WT HSC to MPP Intraphenotypic comparison.

**Table S6**. Differentially expressed genes identified by AHR KO HSC to MPP Intraphenotypic comparison.

**Table S7**. Canonical pathways analysis of the WT intraphenotypic HSC to MPP comparison.

**Table S8**. Canonical pathways analysis of the AHR KO intraphenotypic HSC to MPP comparison.

