## Supplemental Table 1 for "Impact of the aryl hydrocarbon receptor on Aurora A kinase and the G2/M phase pathway in hematopoietic stem and progenitor cells"

| **5’- Primer Sequence** | **Primer name** | **Primer target** |
| --- | --- | --- |
| GTCACTCAGCATTACACTTTCTA | OL4062 | AHR excision forward |
| CAGTGGGAATAAGGCAAGAGTGA | OL4064 | AHR unexcised forward |
| GGTACAAGTGCACATGCCTGC | OL4088 | AHR reverse |
| CAGTGGGAATAAGGCAAGAGTGA | OL4064 | AHR FxFx forward |
| GGTACAAGTGCACATGCCTGC | OL4088 | AHR FxFx reverse |
| CAGTGGGAATAAGGCAAGAGTGA | OL4064 | AHR KO forward |
| AGGGAGATGAAGTATGTGTATGTA | OL4066 | AHR KO reverse |
| AAAGTCGCTCTGAGTTGTTAT | o1MR545 | AHR inducible KO forward |
| GGAGCGGGAGAAATGGATATG | o1MR8546 | AHR inducible KO reverse |
| CCTGATCCTGGCAATTTCG | o1MR8547 | AHR inducible KO Cre specific primer |

**Supplemental Table 1: Primers used for genotyping transgenic mice.**
