## Supplemental Table 2 for "Impact of the aryl hydrocarbon receptor on Aurora A kinase and the G2/M phase pathway in hematopoietic stem and progenitor cells"

| **Marker** | **Fluorochrome** | **Host/ Target** | **Clone** | **Isotype** | **Company** |
| --- | --- | --- | --- | --- | --- |
| CD117 (c-kit) | Brilliant Violet 650 | Rat anti-mouse | 2B8 | IgG2b κ | BD Biosciences |
| CD48 | APC-Cy7 | Hamster Anti-mouse | HM48-1 | IgG1 λ3 | BD Biosciences |
| Lineage Cocktail (CD3/Gr-1/CD11b/CD45R/Ter-119) | Pacific Blue | Anti-mouse | Many | Multiple | Biolegend |
| Lineage Cocktail (CD3/Gr-1/CD11b/CD45R/Ter-119) | eFluor 450 | Anti-mouse | Many | Multiple | Invitrogen |
| CD150 (SLAM) | PE-Cy7 | Rat anti-mouse | TC15-12F12.2 | IgG2a λ | Biolegend |
| CD34 | FITC | Rat anti-mouse | RAM34 | IgG2a κ | BD Biosciences |
| Sca-1 (Ly-6A/E) | PE-CF594 | Rat anti-mouse | A2F10.1 | IgG2a κ | BD Biosciences |
| FLT-3 (CD135) | PerCP-eFluor710 | Rat anti-mouse | A2F10.1 | IgG2a κ | BD Biosciences |
| Aurora A kinase | -- | Rabbit anti-mouse | ST46-07 | IgG |  |
| Anti-rabbit IgG | Alexa Fluor 647 | Donkey anti-rabbit | Poly4604 | Poly Ig | Biolegend |
| Anti-BrdU | Alexa Fluor 647 | Mouse anti-BrdU | 3D4 | IgG1 κ | Biolegend |
| CD117 (c-kit) | APC | Anti-mouse | ACK2 | IgG2b κ | ebioscience |

**Supplemental Table 2. List of antibodies used in flow cytometric analysis of bone marrow cells.**
